# Bimanual digit training improves right-hand dexterity in older adults by reactivating declined ipsilateral motor-cortical inhibition

**DOI:** 10.1101/2021.05.21.445083

**Authors:** Eiichi Naito, Tomoyo Morita, Satoshi Hirose, Nodoka Kimura, Hideya Okamoto, Chikako Kamimukai, Minoru Asada

## Abstract

Improving deteriorated sensorimotor functions in older individuals is a social necessity in a super-aging society. Previous studies suggested that the declined interhemispheric sensorimotor inhibition observed in older adults is associated with their deteriorated hand/finger dexterity. Here, we examined whether bimanual digit exercises, which can train the interhemispheric inhibitory system, improve deteriorated hand/finger dexterity in older adults. Forty-eight healthy, right-handed, older adults (65-78 years old) were divided into two groups, i.e., the bimanual (BM) digit training and right-hand (RH) training groups, and intensive daily training was performed for 2 months. Before and after the training, we evaluated individual right hand/finger dexterity using a peg task, and the individual state of interhemispheric sensorimotor inhibition by analyzing ipsilateral sensorimotor deactivation via functional magnetic resonance imaging when participants experienced a kinesthetic illusory movement of the right-hand without performing any motor tasks. Before training, the degree of reduction/loss of ipsilateral motor-cortical deactivation was associated with dexterity deterioration. After training, the dexterity improved only in the BM group, and the dexterity improvement was correlated with reduction in ipsilateral motor-cortical activity. The capability of the brain to inhibit ipsilateral motor-cortical activity during a simple right-hand sensory-motor task is tightly related to right-hand dexterity in older adults.

## Introduction

Sensorimotor functions deteriorate in older adults [1,2]. Thus, improving these functions in older adults is a social necessity in any super-aging society. Hand/finger dexterity is a representative sensorimotor function that enables the performance of daily skillful manual behaviors. Among primates, humans have well-developed hand/finger dexterity, in which the primary motor cortex (M1) plays a particularly important role [3–6]. Similar to many sensorimotor functions, human hand/finger dexterity is also deteriorated by aging [7–9]. Therefore, improving deteriorated hand/finger dexterity in older individuals is a worthwhile challenge.

Several studies have reported that reduced transcallosal inhibition from the contralateral to the ipsilateral M1 [7] and hyperactivation of the ipsilateral primary sensorimotor cortex (SM1) in older adults [8] are associated with deteriorated dexterity of their right hand, as evaluated by peg task performance. In addition, the aging-related reduction of interhemispheric inhibition exerted from the contralateral (left) M1 may have contributed to the age-related reduction/loss of ipsilateral (right) M1 deactivation during a right-hand sensory-motor task [10]. Thus, these lines of evidence suggest that the reduction/loss of ipsilateral M1 deactivation during a right-hand sensory-motor task, likely caused by the reduction of interhemispheric inhibition from the left to right M1, deteriorates right hand/finger dexterity in right-handed older individuals.

In the present study, first, we confirmed the deterioration of right hand/finger dexterity using a peg task and the reduction/loss of ipsilateral deactivation using a right-hand sensory-motor task (independent from the peg task) in 48 healthy, right-handed, older adults (65–78 years old) compared with 31 younger adults (20–27 years old). Kinesthetic illusion was used as a right-hand sensory-motor task. We measured brain activity using functional magnetic resonance imaging (fMRI) while the blindfolded participants experienced kinesthetic illusory flexion of the right stationary hand elicited by a muscle afferent input during its tendon vibration [11]. It is known that, during the illusion, the hand/arm sections of the contralateral (left) SM1 and of the dorsal premotor cortex (PMD) are activated by receiving the kinesthetic input passively, while the ipsilateral side is deactivated [11], presumably via interhemispheric inhibition from the left to right SM1-PMD. One of the advantages of this task is that it allows the evaluation of the pure state of the interhemispheric inhibitory system by measuring ipsilateral SM1-PMD deactivation (negative blood oxygenation level-dependent [BOLD] signal) in individual participants, with no motor tasks being performed.

Next, prompted by previous reports [7,8], we examined whether the reduction/loss of ipsilateral M1 deactivation observed during the illusion is associated with dexterity deterioration in the older participants. Here, we formulated a specific anatomical hypothesis for the M1 region. Our recent developmental fMRI study showed that deactivation in a particular region of the hand/arm section of the ipsilateral (right) M1 (peak coordinates x, y, z = 36, −26, 66) during a simple right-hand sensory-motor task is better developed in children with higher right hand/finger dexterity [6]. In the present study, we hypothesized that this relationship between deactivation and dexterity observed during childhood is also present in older adults in this particular region. To test this hypothesis, we set a region-of-interest (ROI) in this M1 region (M1 ROI; 4-mm radius sphere around the M1 peak).

Given that the reduction/loss of ipsilateral M1 deactivation (inhibition) is one of the causes of the deterioration of hand/finger dexterity in older adults, their hand/finger dexterity might be improved by reactivating the ipsilateral inhibition, i.e., via interhemispheric inhibition from the left to right M1. Therefore, intensive daily training was carried out by the older participants for approximately 2 months (see Supplementary Methods and Tables S1–3), during which we checked the progress of their training on a weekly basis. The older participants were divided into the following two groups. One group performed bimanual digit training (BM group; n = 23; see Table S1). Bimanual training was chosen because it likely facilitates transcallosal neuronal communication between the two M1 regions [12,13]. During this training, the participants performed both the same and different finger actions using the two hands. In the case of different actions, each of the left or right M1 had to control disparate actions simultaneously while mutually inhibiting the potential occurrence of synchronized actions [14], probably through the transcallosal inhibitory system. Another group underwent unimanual digit training using the right-hand (RH group; n = 25; see Table S1). This group performed the same exercises performed by the BM group but used the right-hand exclusively. Thus, this group intensively trained the right-hand only, and we expected that the interhemispheric inhibition could be better trained in the BM group. Importantly, neither group was trained in the peg task per se during the training period, to avoid peg-task-specific training effects.

After the training, using the peg task, we examined whether the right hand/finger dexterity was improved in the BM group, and whether such behavioral improvement was correlated with the reduction of activity in the M1 ROI (see above) during the illusion. In the series of fMRI results, we also reported the data obtained from whole-brain analysis.

## Results

### Before training

#### Peg task performance

All of the participants performed a 12-hole peg task [6], during which the time required to flip all 12 pegs in each of three trials was measured. Although the average peg time was significantly longer in older adults than in younger adults (22.13 ± 3.65 s [mean ± standard deviation] vs. 20.05 ± 3.19 s; *t*[77] = 2.59, *P* = 0.01), no significant difference in peg time was observed between the two training groups of older adults (BM, 22.23 ± 3.58 s; RH, 22.04 ± 3.78 s; *t*[46] = 0.18, *P* = 0.86; Figure 1, left panel). This indicated that right-hand dexterity was deteriorated in older adults compared with younger adults, and that the pre-training peg time was highly similar between the two groups of older individuals.

**Figure 1.**
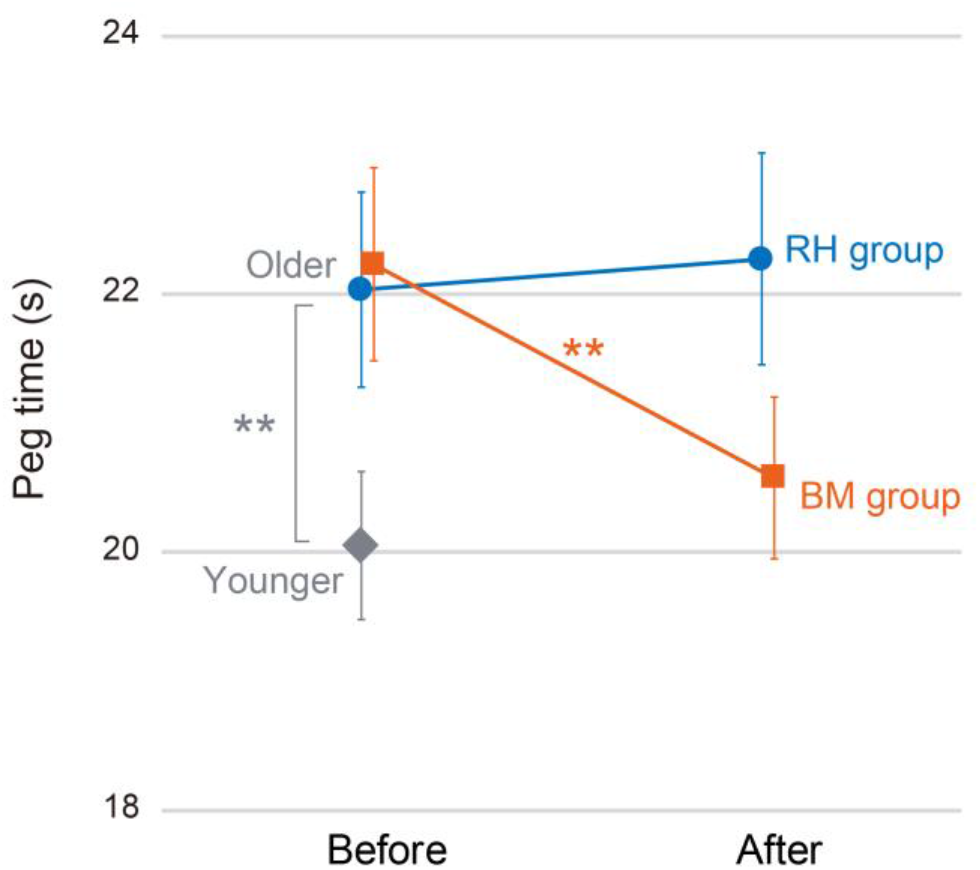
Peg time before (left) and after (right) the training. The orange squares indicate the average peg time in the BM group, the blue circles indicate that obtained in the RH group, and the gray diamond indicates that observed in the younger group. The error bars indicate the SEM. ** *P* ≤ 0.01. Abbreviations: BM, bimanual; RH, right hand; SEM, standard error of the mean.

#### Reduction/loss of ipsilateral deactivation in older adults

Task-related deactivation was examined during the illusion in the entire brain (family-wise error rate [FWE]-corrected extent threshold of *P* < 0.05 across the entire brain for a voxel-cluster image generated at an uncorrected height threshold of *P* < 0.005; blue sections in Figure 2a). We found significant deactivation in the ipsilateral (right) SM1-PMD in the younger group, which was not observed in the older group (Figure 2a; see Table S4a–d for other deactivations and activations).

**Figure 2.**
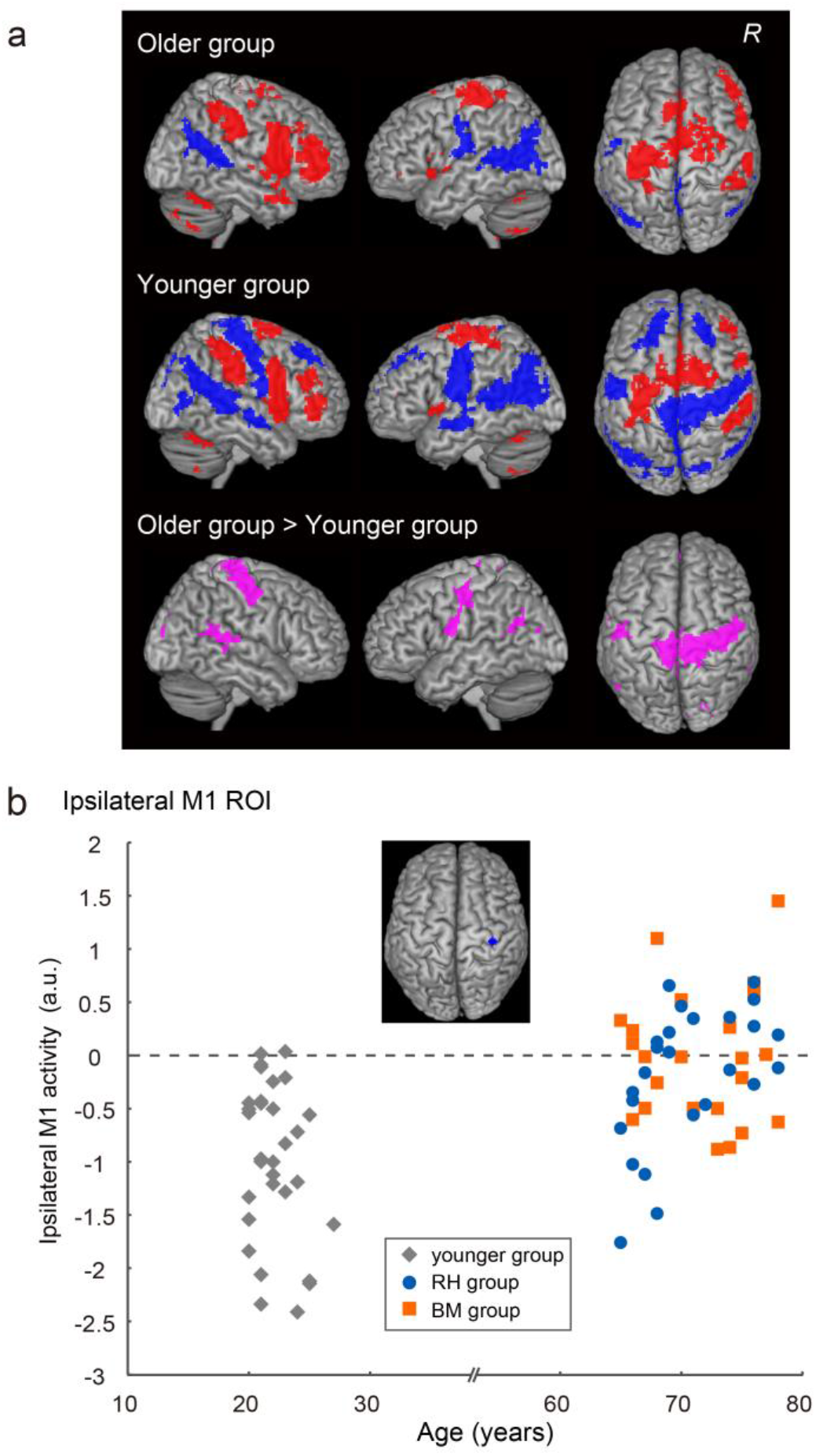
Brain deactivations and activations in the younger and older groups before training. **a**: Brain deactivations (blue) and activations (red) observed during the illusion in the older (top) and younger (middle) groups. Table S4 summarizes the deactivations and activations. The pink areas indicate brain regions with greater activity in the older group than in the younger group (bottom). Table S5 summarizes the age-group difference results, which are rendered on the cortical surface of the MNI standard brain. In each row, the left panel indicates the right-side view, the middle panel indicates the left-side view, and the right panel indicates the top view of the brain. **b**: Relationship between the effect size of ipsilateral M1 ROI activity (a.u.; vertical axis) and age (horizontal axis). The blue section in the top panel indicates the ipsilateral M1 ROI. The data indicated below a dashed line indicate deactivation. The gray diamonds represent the data obtained from the younger group. The orange squares represent individual data obtained from the BM group. The blue circles indicate data from the RH group. Abbreviations: a.u., arbitrary unit; BM, bimanual; MNI, Montreal Neurological Institute; R, right hemisphere; RH, right hand.

The direct comparison of the activity between the two age groups (older vs. younger) revealed several significant clusters of voxels with between-age-group differences in the whole brain (pink section in Figure 2a, bottom panel). The largest cluster was identified in the ipsilateral SM1-PMD and its peak was located in area 4p (see Table S5a for other clusters; see Table S5b for the opposite contrast [younger vs. older]). The ipsilateral SM1-PMD cluster likely covered the hand/arm, trunk, face, and foot sections and extended into the foot section of the contralateral SM1.

We extracted the effect size of brain activity from the ipsilateral M1 ROI (blue section in Figure 2b) in each participant, and plotted it against their individual age. Almost all of the younger adults showed deactivation, whereas deactivation was generally reduced or lost in the older adults (Figure 2b). This finding was corroborated by the analysis of the temporal profile of task-related brain activity during the illusion in a broader region of the hand/arm section of the bilateral SM1-PMD (see Supplementary Results and Figure S1). However, no significant difference was observed between the two training (BM and RH) groups of older adults (*t*[46] = 0.91, *P* = 0.37; Figure 2b). These results were observed even when the two age groups reported almost the same illusory flexion angle of the right-hand (older, 32.50° ± 15.98°; younger, 32.82° ± 13.37°; *t*[77] = 0.09, *P* = 0.93; Table S6). In addition, the BM and RH groups reported almost the same illusory angle before training (*t*[46] = 1.20, *P* = 0.23; Table S6).

#### Correlation between deteriorated dexterity and reduction/loss of ipsilateral deactivation in older adults

Next, we examined whether the reduction/loss of ipsilateral M1 deactivation observed during the illusion is associated with the deterioration of dexterity in the older participants. We performed a regression analysis to depict brain regions in which deactivation was degraded with a longer peg time, i.e., a measure of deteriorated dexterity [7,8]. No significant clusters were found in the whole brain. However, as hypothesized, we found a significant cluster (pink section in Figure 3; peak in area 4a; height threshold *P* < 0.005 uncorrected) in the M1 ROI (blue and pink sections in Figure 3) with small volume correction (SVC; 6 voxels, *P* < 0.05). We also performed the same analysis in the young adults and found no significant clusters in the M1 ROI with SVC. Thus, the correlation between the reduction/loss of deactivation in the focal region of the ipsilateral M1 (M1 ROI) and the deteriorated right-hand dexterity was only observed in the older group.

**Figure 3.**
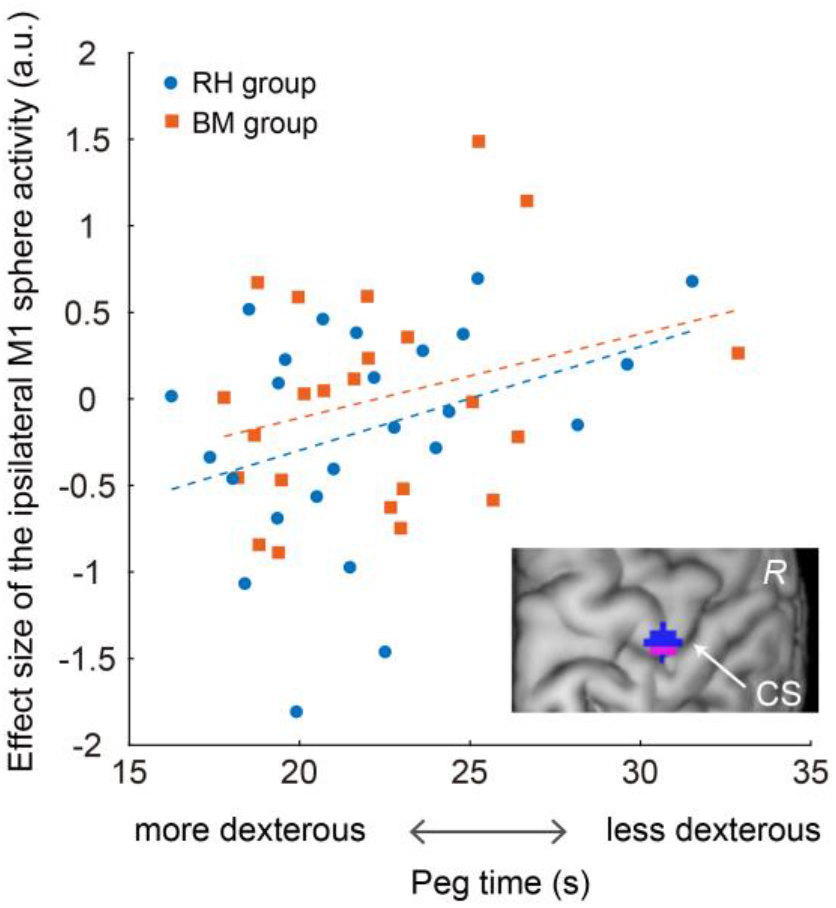
Relationship between the effect size of ipsilateral M1 activity (a.u.; vertical axis) and the peg time (horizontal axis) across 48 older adults. The pink area in the right bottom panel indicates the ipsilateral M1 cluster in which the activity was positively correlated with the peg time. The effect size was extracted from the M1 ROI (blue and pink areas in the right bottom panel). The orange squares represent individual data obtained from the BM group, and the orange dashed line indicates a regression line fitted to the data. The blue dots indicate individual data from the RH group, and the blue dashed line indicates a regression line fitted to the data. Abbreviations: a.u., arbitrary unit; BM, bimanual; CS, central sulcus; R, right hemisphere; RH, right hand.

Such a positive correlation in the older group was confirmed by visualizing the relationship between the activity obtained from the M1 ROI and the peg time (*r* = 0.32, n = 48, *P* < 0.05; Figure 3). As shown in Figure 3, in the older adults, a greater reduction or loss of ipsilateral M1 deactivation was associated with a greater deterioration of their right-hand dexterity. Importantly, no significant between-training-group differences were observed either in the slopes of the regression lines fitted to the data obtained from each group (BM or RH; F[1, 44] = 0.054, *P* = 0.82) or in their intercepts (F[1, 45] = 0.859, *P* = 0.36), indicating that the two regression lines were not different. Thus, the results indicate that activity in the ipsilateral M1 region is related to dexterity deterioration in the older adults. The results reported above were observed in the absence of correlations between the illusory angle and the peg time (*r* = 0.11, n = 48, *P* = 0.45) and between the angle and the ipsilateral M1 ROI activity (*r* = 0.15, n = 48, *P* = 0.32) among the older participants.

### After training

#### Peg task performance

We found no significant between-training-group differences in the number of training sets performed by each group (see Supplementary Methods and Table S3). Despite the absence of significant differences, we observed peg time improvement only in the BM group (Figure 1). In this group, the peg time improved from 22.23 ± 3.58 s (before) to 20.58 ± 3.00 s (after). A two-way mixed-design analysis of variance (ANOVA; group [BM and RH] × order [before and after]) of peg time showed a significant interaction between group and training (F[1, 46] = 7.67, *P* < 0.01). A post hoc analysis revealed that the peg time became significantly shorter in the BM group after training (*t*[22] = 3.70, *P* < 0.005 after Bonferroni correction), whereas no significant change was observed in the RH group (*t*[24] = −0.46, *P* > 1 after Bonferroni correction). These results showed that bimanual training, but not unimanual RH training, improved right hand/finger dexterity effectively.

#### Correlation between dexterity improvement and reduction of ipsilateral M1 activity

Because we found an improvement in dexterity only in the BM group, we performed a regression analysis to examine if the peg time improvement was associated with activity reduction after the training (after–before) in this group. We found a significant cluster in the hand/arm section of the ipsilateral SM1-PMD (Figure 4a), in addition to a cluster in the hand/arm sections of the bilateral caudal cingulate motor areas in the entire brain (see Table S7). Importantly, the ipsilateral SM1-PMD cluster overlapped with the M1 ROI (white section in Figure 4a), in which we found a correlation between the degree of reduction/loss of deactivation and the deterioration of dexterity before training (blue and pink sections in Figure 3). In contrast, in the RH group, we found no significant clusters with such a correlation in the entire brain.

**Figure 4.**
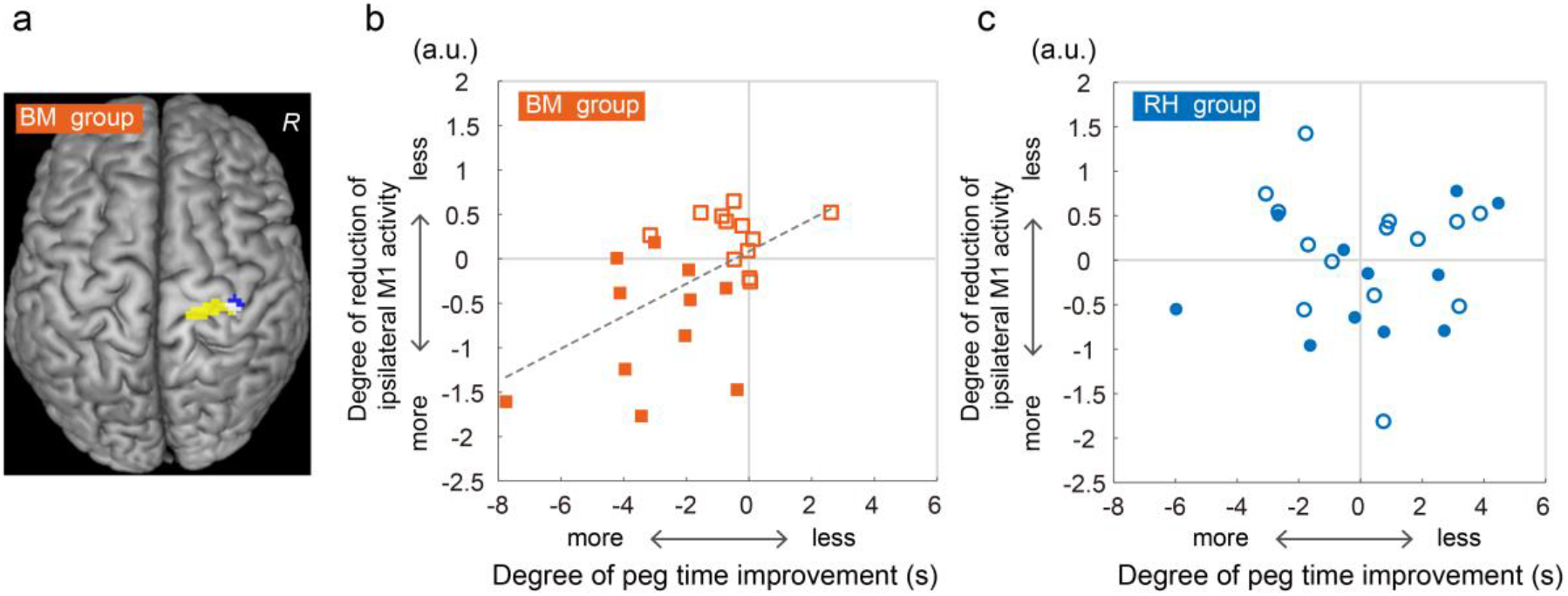
Relationship between the degree of reduction in ipsilateral M1 activity and peg time improvement. **a**: Significant cluster (yellow and white areas) in which the degree of activity reduction was correlated with peg time improvement in the BM group. The cluster is superimposed on the MNI brain. **b**, **c**: Relationship between the degree of reduction of activity (a.u.; vertical axis) in the M1 ROI (blue and white areas in panel A) and peg time improvement (s; horizontal axis) in each group (b for the BM group, c for the RH group). In each panel, the filled symbols represent data obtained from the participants who exhibited ipsilateral M1 hyperactivation before the training (ipsilateral M1 showed activity > 0). The open symbols represent data obtained from the participants in whom ipsilateral M1 deactivation was preserved before the training. The dashed line in panel B indicates a regression line fitted to the data. Abbreviations: a.u., arbitrary unit; BM, bimanual; MNI, Montreal Neurological Institute; R, right hemisphere; RH, right hand.

Finally, we plotted the degree of reduction of activity (after–before) obtained from the M1 ROI against the peg time improvement (after–before) in each group, and confirmed the presence of a positive correlation in the BM group (*r* = 0.54, n = 23, *P* = 0.008; Figure 4b). This implies that participants with a greater peg time improvement had a greater reduction of ipsilateral motor-cortical activity after the training. This correlation was observed even though there were no correlations between the change in illusory angle (after–before) and the peg time improvement (*r* = 0.12, n = 23, *P* = 0.59) and between the change in angle and the degree of activity reduction (*r* = 0.26, n = 23, *P* = 0.23) among the participants in the BM group. The RH group showed no such correlation (*r* = 0.03, n = 25, *P* = 0.86; Figure 4c).

Notably, in the BM group, the peg time improvement and the degree of reduction in ipsilateral M1 ROI activity were significantly greater in participants with ipsilateral M1 hyperactivation (greater than 0) before training (n = 11; filled squares in Figure 4b) compared with those with ipsilateral deactivation (n = 12; open squares in Figure 4b; *t*[21] = 3.73, *P* < 0.005 for improvement, and *t*[21] = 4.52, *P* < 0.001 for reduction after Bonferroni correction). These findings were observed even when the number of training sets did not differ between these two subgroups (*t*[21] = 1.54, *P* = 0.14). In contrast, no such differences were observed in the RH group (*t*[23] = 2.07, *P* > 1 for improvement; *t*[23] = 1.04, *P* = 0.62 for reduction after Bonferroni correction; Figure 4c). Hence, the bimanual training resulted in significantly greater behavioral and neuronal training effects in participants who had ipsilateral M1 activation (lost ipsilateral deactivation) before the training.

## Discussion

This study clearly demonstrated that ipsilateral motor-cortical activity as an index of declined interhemispheric inhibition during a simple right-hand sensory-motor task was associated with deterioration of right hand/finger dexterity in older individuals, and that bimanual digit training, which is available for anyone at any time and in any place, can improve unimanual dexterity by causing a reduction of ipsilateral motor-cortical activity. These results revealed a tight relationship between interhemispheric inhibition and hand/finger dexterity in older adults, and suggest the trainability of the interhemispheric inhibitory system, which is deteriorated during normal aging, through bimanual training.

### Physiological and technical considerations

In younger adults, ipsilateral SM1-PMD deactivation has been frequently reported during right-hand motor and proprioceptive tasks [15–18]. The physiological mechanisms underlying the task-induced negative BOLD phenomenon, which is indicative of deactivation, are not fully understood [19,20]. However, many recent studies have suggested that this phenomenon is associated with neuronal inhibition (see the discussion in reference 21) [21], with no exception in the cerebro-cerebellar sensorimotor network [22,23].

The reduced interhemispheric inhibition exerted from the left M1 has been shown to be associated with a reduction/loss of deactivation in the right M1 in older adults [10,24], and that the interhemispheric (transcallosal) inhibition exerted from the M1 to its opposite M1 can suppress activity in the latter [25–28]. Thus, the ipsilateral M1 deactivation observed during the right-hand illusion might be derived, at least partly, from interhemispheric inhibition exerted from the hand/arm section of the contralateral (left) M1, which was activated during the illusion (Figure 2a), although we cannot fully exclude the possibility of inhibition from other brain structures, such as the thalamocortical system [29], the PMD [30,31] and SMA [32].

In the current work, we evaluated ipsilateral sensorimotor deactivation (inhibition) using a proprioceptive task independent from the peg task, because such a complex finger task might activate the ipsilateral SM1-PMD, even in younger adults [33], thus precluding the proper evaluation of the entity of ipsilateral inhibition in individual brains if brain activity is scanned using the peg task. In addition, ipsilateral SM1-PMD activity (i.e., reduction/loss of deactivation) observed in the older adults during the non-motor task indicates that this phenomenon is not related to compensation of motor functions by the ipsilateral SM1-PMD (see below).

It has been suggested that, in older adults, a higher level of performance in a motor task is related to a higher capacity to modulate motor-cortical inhibition when performing the task [34], which largely depends on its basic state of local inhibition [35]. We assumed that the ipsilateral SM1-PMD deactivation observed during the illusion may reflect the “basic state” of its local inhibition, because this is likely caused mainly by the interhemispheric inhibition exerted from the contralateral SM1-PMD when this region is activated by simply receiving a kinesthetic input. Therefore, the reduction/loss of ipsilateral SM1-PMD deactivation (i.e., basic state) observed during the illusion might be associated with a reduced capability to modulate ipsilateral motor-cortical inhibition when the older participants performed the peg task; however, the confirmation of this claim should be the subject of future studies.

### Deteriorated dexterity and its relation to ipsilateral activity in older adults before training

The degree of reduction/loss of ipsilateral M1 deactivation was correlated with deteriorated dexterity in the older group (Figure 3). This finding was confirmed by the analysis performed using a larger sample of older participants (see Supplementary Results and Figure S2). Thus, these findings are consistent with those reported previously [7,8]. Notably, the M1 region is identical to the region that was identified in our previous study, which showed that the degree of deactivation was associated with a better right hand/finger dexterity during childhood [6]. Interestingly, such a correlation was not observed in the younger group, as demonstrated in our previous study [6]. In another study, we also demonstrated that ipsilateral M1 deactivation during a right-hand unimanual motor task progresses from childhood to adolescence, but stabilizes from adolescence to adulthood [21], indicating that the functional differentiation between the left and right M1 occurs between childhood and adolescence. Therefore, ipsilateral M1 deactivation might be deeply associated with hand/finger dexterity during childhood, when the deactivation (inhibition) is developing, and during elderhood, when it deteriorates. These findings suggest that ipsilateral M1 inhibition in the central motor system substantially influences dexterous motor control during childhood and old age.

In nonhuman primates, motor neurons in lamina IX of the spinal cord directly receive efferents primarily from the contralateral M1. This indicates that the fast, direct corticomotor pathway, which enables fine, dexterous finger movements [4,5], originates primarily from the contralateral M1, particularly the new M1 (the caudal region of the M1) [36], which likely corresponds to the human area 4p [37]. Although the ipsilateral M1 also projects to the spinal cord, there are fewer terminals in lamina IX [38], indicating that the ipsilateral M1 projection is not suitable for the direct, fine control of hand/finger muscles; rather, it is suitable for muscle synergy control. The development of ipsilateral M1 deactivation that occurs from childhood to adolescence [21], together with the development of transcallosal inhibition during these periods [39], suggests that the brain develops to facilitate dominance of the contralateral M1 for the fine control of dexterous hand/finger movements by inhibiting the ipsilateral M1, although the brain is capable of recruiting the ipsilateral M1 in cases of brain stroke [40–42]. We assumed that the capability to inhibit the ipsilateral M1 allows the brain to avoid potential disturbance from the M1, for elaborate control by the contralateral one [6]. This view seems to be compatible with our findings that ipsilateral M1 activity was correlated with deteriorated dexterity (Figure 3) and that the reduction of ipsilateral M1 activity was associated with dexterity improvement (Figure 4) in older adults, who may partially utilize the ipsilateral M1 when performing a motor task [43].

We assumed that a possible decline in interhemispheric inhibitory function, which can suppress and regulate ipsilateral M1 activity (see above), is one of the factors that contribute to the reduction/loss of ipsilateral M1 deactivation (inhibition) in older adults. Although little is known about the exact neuronal mechanisms underlying decreased interhemispheric inhibition, the quantitative and qualitative deterioration of transcallosal nerve fibers observed in older adults could be related to this decline [13,44]. Moreover, a reduction in the level of the inhibitory neurotransmitter gamma-aminobutyric acid (GABA) in the frontoparietal cortices of older adults must also be deeply correlated with this phenomenon [45].

If we consider the relationship discovered here between the reduction/loss of ipsilateral M1 deactivation and the deteriorated dexterity observed in the healthy older adults (Figure 3) together with the fact that patients with stroke who lack dexterity often exhibit ipsilateral SM1-PMD activity, we may assume that ipsilateral SM1-PMD activity is an index of the deterioration of dexterity in these individuals. This view does not contradict the fact that the compensatory ipsilateral SM1-PMD activity that occurs after contralateral SM1-PMD stroke often disappears when hand/finger dexterity improves [41]. The reduction/loss of ipsilateral M1 deactivation detected during the illusion in the older adults (Figure 2b) likely represents a chronic decline in ipsilateral M1 inhibition. As described above, ipsilateral SM1-PMD activity is often reported when people perform complex finger movements [33] and precision-demanding tasks [46,47]. This could be caused by an “active” disinhibition, in which the brain utilizes the ipsilateral SM1-PMD by affirmatively releasing interhemispheric inhibition; this could be distinct from the chronic decline observed in older adults. In turn, this decline in older adults might indicate a lesser potential of their brains to disinhibit the ipsilateral M1 (i.e., compensation by the ipsilateral M1) in cases of stroke.

### Improvement of right-hand dexterity and its relation to reduced ipsilateral M1 activity after training

Bimanual training improved right-hand dexterity (Figure 1) in association with reduced ipsilateral M1 activity during the illusion (Figure 4b). Because the present training did not include the peg task training per se, we attributed this behavioral improvement to non-peg-task-specific neuronal changes caused by the bimanual training, i.e., most likely the reactivation of interhemispheric inhibition. Importantly, the bimanual training resulted in significantly greater behavioral and neuronal training effects in participants who exhibited ipsilateral hyperactivation before the training (Figure 4b), clearly indicating the effectiveness of bimanual training for older adults who need reactivation of ipsilateral inhibition.

The claim that bimanual training reactivates interhemispheric inhibition seems to be supported by the following evidence. The analysis of functional connectivity during the illusion revealed that the bimanual training reduced the functional connectivity between the bilateral SM1-PMDs compared with the RH training (see Supplementary Results, Figure S3, and Table S8). In the present study, no significant correlation was observed between the reduction of interhemispheric functional connectivity and dexterity improvement. However, a previous study reported that the greater functional connectivity detected between the left and right M1 regions in older adults, which might reflect an aging-related release from the normally predominantly inhibitory interhemispheric communication, was associated with poorer performance in an unimanual finger tapping task [12]. The decrease in interhemispheric connectivity observed here after the bimanual training suggests that the brain changed to afford the activation of the two SM1-PMDs independently of each other. This suggests that the transcallosal inhibitory system is reactivated after the training to suppress a more synchronized activity between the two SM1-PMDs before the training. Thus, the various findings reported here indicate that bimanual training may reactivate the interhemispheric inhibitory system between the bilateral SM1-PMDs more effectively than unimanual RH training. However, the lack of electrophysiological measurement of interhemispheric inhibition using transcranial magnetic stimulation was one of the limitations of the present study. Therefore, further studies are needed to support our claim. However, if we consider the study that reported that 12-week aerobic whole-body exercises improved interhemispheric inhibition and hand/finger dexterity in older adults [48], then the interhemispheric inhibitory system, which deteriorates with normal aging, can be trained.

Theoretically, the RH training could also lead to an inhibitory effect on the ipsilateral M1, because it has been shown that, in younger adults, the exclusive use of one hand elevates the motor-cortical excitability in the contralateral M1, which leads to a stronger inhibitory effect on the ipsilateral M1 [49]. However, our results suggest that, in older adults in whom transcallosal inhibition is likely deteriorated [7,12], unimanual training might not effectively decrease ipsilateral M1 activity, probably because of the coactive mode between the bilateral SM1-PMDs (see Figure S1).

### Ipsilateral M1 activity and hemispheric asymmetry reduction in older adults (HAROLD)

Our finding that ipsilateral M1 activity, as an index of declined interhemispheric inhibition, was associated with the deterioration of right-hand dexterity in older adults was compatible with previous reports [7–9,48]. Nevertheless, the finding that a faster reaction time in a simple button-press task was correlated with greater ipsilateral premotor activity in older adults is contradictory [50]. It remains controversial whether additional activation and overactivation in older adults reflect compensatory recruitment or reorganization [1] or are simply a consequence of an age-related decline in the inhibitory process [51]. For example, it seems that the additional activations in the left frontoparietal cortices (but not in the ipsilateral SM1-PMD) that occur during a right-hand–foot coordination task can be considered as the former [52], whereas declined interhemispheric inhibition can be considered as the latter. The latter view seems to also be supported by reports that motor overflow (involuntary movement or muscle activity) in passive homologous muscles contralateral to voluntary movement is often greater in older adults vs. younger adults, possibly because of a decline in interhemispheric inhibition in older adults [53].

Prefrontal activity during cognitive (e.g., memory) tasks tends to be less lateralized (i.e., showing bilateral activity) in older adults vs. younger adults. This phenomenon is called hemispheric asymmetry reduction in older adults (HAROLD) [54]. The reduction/loss of ipsilateral SM1-PMD deactivation observed during the illusion in older adults seems to be a type of HAROLD (Figure 2b). The present sensorimotor HAROLD (i.e., reduction/loss of ipsilateral M1 deactivation) was associated with deteriorated dexterity (Figure 3; see also references[7–9,48]), whereas prefrontal HAROLD may have a compensatory function for maintaining cognitive performance [55]. A study of resting-state functional connectivity has suggested that cross-hemispheric connectivity is higher in the sensorimotor cortex, whereas within-hemispheric connectivity is higher in the prefrontal cortex, in younger adults [56]. Thus, interhemispheric inhibition between two sensorimotor cortices might be necessary for the brain to utilize each hemisphere independently against their intrinsically synchronous mode. Hence, we raise the possibility that HAROLD in the bilateral primary sensorimotor cortices can be observed as a mere result of the age-related decline of interhemispheric inhibition, as if the system becomes infantilized, although the sensorimotor HAROLD may become beneficial when the brain generates symmetrical bimanual movements by recruiting the bilateral sensorimotor cortices in a coactive mode (see also reference [53]), as well as in cases that require a faster reaction that does not require dexterity [50].

The brain of older individuals can be characterized by a reduction/loss of various interregional brain deactivations (inhibition) during cognitive and sensorimotor tasks [57–60]. The current work suggests that training tasks that can facilitate interregional brain communications might improve the cognitive and sensorimotor functions that are deteriorated in older individuals by reactivating their declined interregional inhibitory system that prevails in younger adults.

## Methods

### Participants

Healthy, right-handed younger adults (n = 31; age, 22.1 ± 1.8 [mean ± standard deviation] years; age range, 20–27 years; males, 22) and older adults (n = 48; age, 65–78 years) participated in this study. We divided the older participants into two training groups: the RH group (n = 25; age, 70.6 ± 4.2 years; males, 17) and the BM group (n = 23; age, 71.7 ± 4.3 years; males, 14). This was a single-blinded, randomized, controlled study (see Supplementary Methods). We assessed the cognitive status of older participants using the Mini-Mental State Examination. All participants scored higher than the cut-off score of 24 [61], and no significant difference was observed between the RH (28.4 ± 1.9) and BM (28.8 ± 1.6) groups. We confirmed the handedness of the participants using the Edinburgh Handedness Inventory [62]. No participants had a history of neurological, psychiatric, or movement disorders based on self-reports.

The study protocol was approved by the Ethics Committee of the National Institute of Information and Communications Technology. We explained the details of the present study to all participants before the experiment, and they then provided written informed consent. The study was conducted according to the principles and guidelines of the Declaration of Helsinki (1975).

### Peg task to evaluate hand/finger dexterity

We used a 12-hole peg task to evaluate right-hand/finger dexterity [6], because peg tasks have generally been used to evaluate hand/finger dexterity, particularly that of the fingertips, across individuals with a wide age range [7–9]. Participants had to remove a small peg that had been inserted in one of 12 holes on a board using their right fingers, vertically flip the peg, and reinsert the peg into the same hole, repeatedly. We measured the time required to flip all 12 pegs using a stopwatch. Because none of our participants had previously experienced this task, each participant performed the peg task three times. In this task, a participant who is better able to coordinate her/his fingertip movements rapidly, without generating superfluous movements, should be able to complete the task more quickly.

Before training, we calculated the average time (peg time) of three trials in the older and younger groups, respectively. We evaluated the between-age-group differences by conducting a two-sample *t*-test. To evaluate the training effect on peg task performance, we also calculated the peg time for the two training groups after training. We conducted an ANOVA that included one between-subject factor (group [2]: BM or RH) and one within-subject factor (order [2]: before and after), and further performed a post hoc test with Bonferroni correction to evaluate the possible training effects in each group.

### Two-month training

Only the older participants performed approximately 2 months of daily training. For each training group, we prepared five types of training menus; the details of the menus performed by each group, the training procedure, and the mean number of sets of training menus performed during the training period are provided in the Supplementary Methods and Tables S1–3. The participants had to perform these menus at home every day (homework) until the day before the second MRI day.

### fMRI task

#### Kinesthetic illusion task

A kinesthetic illusion task was used to evaluate the ipsilateral SM1-PMD deactivation, as described previously [63]. Briefly, we vibrated the tendon of the extensor carpi ulnaris muscle of the relaxed right wrist, which elicited an illusory flexion of the stationary right-hand [11]. Detail of this task procedure is described in the Supplementary Methods.

One run consisted of five tendon-vibration epochs (each lasting for 15 s). The tendon-vibration epochs were separated by 15-s baseline periods. During the baseline period, we vibrated the skin surface over a nearby bone (i.e., the processus styloideus ulnae of the hand next to the tendon; bone vibration) using the same stimulus, which mainly elicits a cutaneous vibration sensation with no reliable illusion, to control skin vibration and attentional effects [63]. Each run also included a 25-s period before the start of the first epoch.

During fMRI scanning, we asked the participants to close their eyes, relax their entire body, refrain from producing unnecessary movements, and be aware of movement sensations from the vibrated hand (during both tendon vibration and bone vibration). After each experimental run, we measured maximum illusory angle that they experienced in a run (see detail in Supplementary Methods), and the mean illusory angle between two runs was calculated for each participant. Detail of statistical analyses are described in the Supplementary Methods. The illusory angles recorded in each group are summarized in Table S6.

#### Single-subject analysis

Details of MRI data acquisition and image preprocessing are described in the Supplementary Methods. After the conventional preprocessing of MRI data, we used a general linear model [64] to analyze fMRI data. We prepared a design matrix for each participant to analyze the functional images before training. The design matrix contained a boxcar function for the task epoch in each run, which was convolved with a canonical hemodynamic response function. To correct for residual motion-related variance after realignment, six realignment parameters were also included in the design matrix as regressors of no interest. In the analysis, we did not perform global mean scaling to avoid inducing type II errors in the assessment of negative BOLD responses [65]. We generated an image showing task-related activation/deactivation in each participant. For older adults, we generated another contrast image showing the difference between the results obtained before and after training (after–before) in each older participant. These images were used in the subsequent second-level group analyses.

#### Analyses before training

We examined task-related activation/deactivation during the illusion using a second-level group analysis

[66] with a one-sample *t*-test to illustrate the patterns of task-related activation/deactivation across the entire brain in each age group (Figure 2a). We also examined between-age-group differences by comparing the older group with the younger group (older vs. younger). We used a family-wise error rate [FWE]-corrected extent threshold of *P* < 0.05 across the entire brain for a voxel-cluster image generated at the uncorrected height threshold of *P* < 0.005, which was consistently used in the present whole-brain analyses. For the anatomical definition of the identified peaks, we referred to the cytoarchitectonic map implemented in the SPM Anatomy toolbox [67], which was also consistently used in the present study. To confirm the between-age-group differences in the ipsilateral (right) M1 region, we set an ipsilateral M1 ROI as a 4-mm radius sphere around the M1 peak (36, −26, 66), in which the degree of deactivation is correlated with a better right hand/finger dexterity during childhood [6]. We extracted the effect size of brain activity from the M1 ROI for each participant and plotted it against their individual ages (Figure 2b).

We performed a regression analysis to test whether the degree of reduction/loss of ipsilateral M1 deactivation is associated with dexterity deterioration in the older group (n = 48). In this analysis, we hypothesized that a region with such a correlation could be observed in the M1 ROI, in which the degree of deactivation is correlated with a better right hand/finger dexterity during childhood [6]. Based on this strong anatomical hypothesis, we applied an SVC (*P* < 0.05) [68] with the M1 ROI (above). Finally, to illustrate the relationship between the individual degree of ipsilateral M1 activity and the individual peg time in the older group, we extracted the data from the M1 ROI and plotted them against the peg time (Figure 3). We fitted a regression line to the data obtained from each group (BM and RH). We used an analysis of covariance for the statistical evaluation of the slopes of the regression lines and their intercepts.

#### Regression analysis between peg time improvement and brain activity change

Because we found that the dexterity improved in the BM group exclusively, we further conducted a regression analysis to examine brain regions in which the change in activity observed after the training was associated with the improvement in peg time in this group. To define the individual peg time change, we subtracted the peg time before the training from that after the training (after–before) for each participant. We used the individual contrast image (after–before) to identify brain regions in which the activity change was correlated with the individual peg time change in the entire brain (Figure 4a). The same analysis was performed for the RH group. To verify the correlation observed in the BM group, we plotted the individual degree of reduction in ipsilateral M1 activity obtained from the M1 ROI (see above) against the peg time improvement in each group (Figures 4b, c). Finally, because we found that the peg time improvement and the degree of reduction in ipsilateral M1 ROI activity in the BM group were greater in participants (n = 11) with ipsilateral hyperactivation in the M1 ROI (> 0) before the training than in those (n = 12) with ipsilateral deactivation, we conducted a post hoc two-sample *t-*test with Bonferroni correction to evaluate possible between-subgroup differences. We performed the same post hoc analysis for the RH group.

## Data availability statement

The datasets generated during and/or analyzed during the current study are available from the corresponding author on reasonable request.

## Acknowledgments

The authors are grateful to Ms. Chie Kawakami and Ms. Keiko Ueyama and the CiNet technical staff for their support during the study. The first author is also grateful to Mizuno Corporation for financial and training support for this study, especially to Mr. Kunio Yoshikawa, Ms. Naoko Matsuo, and Ms. Mikiko Kazama for their support in training, and Mr. Yasunori Kaneko and Mr. Hiroshi Nagao for their useful discussions. We would like to thank enago for English language editing.

## Author Contributions

Author contributions included conception and study design (EN, TM, HO, CH, and MA), data collection or acquisition and statistical analysis (EN, TM, SH, NK, HO, and CK), interpretation of results and drafting the manuscript work or revising it critically for important intellectual content (EN, TM, SH, NK, and MA), and approval of final version to be published and agreement to be accountable for the integrity and accuracy of all aspects of the work (All authors).

