## Supplementary information for "Bimanual digit training improves right-hand dexterity in older adults by reactivating declined ipsilateral motor-cortical inhibition"

Eiichi Naito, PhD

**This file includes:**

Supplementary text

Figures S1 to S3

Tables S1 to S8

SI References

**Supplementary Information Text**

**Supplementary Methods**

#### Participants

The younger participants were recruited from local universities, whereas the older participants were recruited from a local human resources center for the elderly. When recruiting older adults, we advertised two schedules for the MRI date and training period for the RH and BM groups, respectively. Participants chose either schedule according to their convenience, and they were not informed of the group to which they belonged. Thus, this was a single-blinded, randomized, controlled study. Approximately 1 month before the first MRI experiment, we held a briefing session for each group of older participants in which we explained the content of the study. At that time, we did not inform them of the training menus. All participants provided written informed consent. We also explained the details of the MRI experiment to the younger participants before the experiment, and they then provided written informed consent.

#### Two-month training

The training menus are summarized in Table S1. These menus were designed by professional trainers at Mizuno Corporation. The training menus for the BM group were selected from those used in the exercise class for older people (LaLaLa Circuit Lite, https://www.mizuno.jp/facility/facility-event/lalalafit.aspx) conducted by Mizuno Corporation. For the RH group, the menus were arranged such that the exercises could also be performed using one hand.

**Menu 1 Finger extension.** The participants extended one finger at a time from the metacarpophalangeal (MP) joint, from the first to the fifth finger. The exercise was repeated from the fifth to the first finger. Movements from the first to the fifth finger and from the fifth to the first finger were defined as one set. Participants in the RH group performed the exercises using their right hand only while it rested hand on their thighs. The BM group performed the exercises using both hands simultaneously, with the palms together.

**Menu 2 Finger counting.** The participants flexed or extended one finger at a time from the MP joint, counting aloud from 1 to 10. Movements up to a count of 10 were considered one set. Participants in the RH group performed the exercises using their right hand only. Before initiating the exercise, only the first finger of the right hand was flexed, and the exercise was started with the finger in that position. Participants flexed one finger at a time from the MP joint, starting from the second to the fifth finger, while counting aloud from 1 to 4 (red numbers in Table S1). Once all fingers were flexed, they were then extended one finger at a time from the MP joint, beginning from the fifth finger and up to the first finger, while counting aloud from 5 to 9 (red numbers in Table S1). Finally, they flexed the first finger while counting to 10, to complete the exercise. The BM group performed the exercise using both hands. Similar to the RH group, before beginning the exercise, the BM group flexed only the MP joint of the first finger on the right hand and extended all MP joints of the left hand. The right-hand movements were identical to those performed in the RH group; however, the left hand moved different fingers. On the left hand, the participants flexed one finger at a time from the MP joint, beginning from the first finger and up to the fifth finger, while counting aloud from 1 to 5. Subsequently, they extended the flexed fingers, one finger at a time at the MP joint, beginning from the fifth finger and up to the first finger, while counting aloud from 6 to 10. In other words, the BM group performed different finger movements with the right and left hands.

**Menu 3 Finger rotation.** The participants rotated one finger at a time in one direction for approximately 3 s and then rotated it in the opposite direction for approximately 3 s. The order of rotation direction was not stipulated. This movement was performed from the first to the fifth finger and repeated from the fifth finger back to the first finger. Ten finger movements were considered as one set. The RH group performed this exercise using the right hand only, with the palm resting on the thigh. The BM group started with the tips of the fingers of both hands, moved the same finger of both hands in the same direction, and rotated the fingers of both hands simultaneously, with the fingers half a circumference out of synchronization. In other words, the BM group was required to control the movement of the left and right fingers, with the rotation being out of synchronization.

**Menu 4 Rock–paper–scissors (gu-choki-pa).** The participants made the shape of “pa(paper)/choki(scissors)/gu(rock)” with their hands while saying “gu/choki/pa” aloud simultaneously (red words in Table S1). Ten repetitions of the movement were considered as one set. The RH group performed exercises using the right hand only. Therefore, in the RH group, the movement of the right hand did not match the spoken word, with the exception of “choki.” In the BM group, the right-hand movement was identical to that in the RH group; however, the left hand made finger shapes in the same order as the spoken words (gu/choki/pa). Therefore, this group had to repeat the movements with the left hand, and the right-hand movements did not match the spoken words, with the exception of “choki.”

**Menu 5 Finger shape generation.** The participants repeatedly created finger shapes corresponding to each number while saying the numbers 1 to 5 aloud (red numbers in Table S1). Finger shapes were generated by extending the corresponding finger to the MP joint. Participants counted aloud from 1 to 5, followed by 5 to 1, and this combination was considered as one set. Different fingers were extended at different number counts: number 1, only the first finger; 2, the fourth and fifth fingers; 3, the first, second, and third fingers; 4, the second, third, fourth, and fifth fingers; and 5, all fingers. While counting from 1 to 4, movements were performed with the palms facing down; when number 5 was called out, the forearm was supinated to turn the palm upward. The RH group performed this exercise using the right hand only, whereas the BM group performed the movements using both hands simultaneously.

Only the older participants performed approximately 2 months of training. For each group, we held a weekly training session in which all subjects had to participate. We started the first training session after all the participants in each group completed their first MRI experiments. Each weekly training included some manual and fitness training, which was demonstrated and accompanied by a professional instructor. Importantly, in this weekly session, we introduced the training menus and asked the participants to perform the exercises on the menus at home every day, as homework, for the next 6 days; moreover, we taught them how to perform the exercises. We did not particularly ask participants to perform homework on the day of the weekly training session. The participants in a group were strictly prohibited from describing their training menus to those in the other group; thus, the participants in a group did not know the training menus performed by those in the other group. During the training period, participants were also strictly prohibited from practicing the peg task and from starting to learn new motor tasks, particularly those involving the hands.

When we explained the homework during the first weekly training session, we motivated the participants to carry it out by informing them that, if they worked hard to complete their homework, they would receive small gifts on the last week of the experiment. The rewards were a hand towel and an exercise instruction book, in addition to a certificate of training completion.

We did not give out the rewards depending on the amount of homework performed; rather, we gave the same rewards to all participants.

Three menus were selected from the five menus, to ensure that participants did not become bored by always having to follow the same menu, and these were selected as homework menus for each week (Table S2). In the first week, only two types of exercises were selected. The participants were asked to perform at least three or five sets from each menu every day. We decided on the number of sets in accordance with the difficulty of each menu and the number of weeks of training. The participants recorded the number of homework sets that they performed each day on the record sheets provided by the experimenters. In the weekly session, the experimenters checked the participants' weekly training progress based on the record sheets, and encouraged them to perform at least the required number of training sets.

We prepared seven regular training sessions. As the length of motor practice that is needed to induce neuronal changes in the brain is unclear, we decided on a training period based on scientific, empirical, and practical reasons. Scientifically, neuronal changes in the brain have often been reported in younger adults after several (3–6) weeks of motor practice [1]. In the case of older adults, brain activity changes were reported after 6 weeks of motor practice [2]. Empirically, professional trainers at Mizuno Corporation also gained the impression that older adults generally became better able to perform the motor program of the LaLaLa Circuit Lite after 4–6 weeks of this practice. Therefore, we assumed that a training schedule of more than 6 weeks was necessary to induce behavioral and neuronal changes in older adults. Practically, a 7-week training period was our limitation in terms of the use of facilities for weekly sessions and recruiting professional instructors. Thus, even though we found both neuronal and behavioral changes after approximately 2 months of training in the present study, we are uncertain whether a 2-month training is necessary and whether the training effects can be observed using a shorter period of training.

During the regular training weeks, five participants in the BM group and five participants in the RH group missed one day of the weekly training sessions. For those participants, we prepared an additional week, so that all of the participants completed the 7 weeks of regular training. We started the second MRI scan during the $8^{th}$ week of the experiment. Because the second MRI days varied across participants, we asked them to continue the homework of the $8^{th}$ week and to record it until the day before the second MRI day. The participants handed their recording sheets to the experimenters on their second MRI day. The average intervals between the first training day and the second MRI experiment in each group were 54.4 ± 3.5 days for the BM group and 55.4 ± 2.7 days for the RH group.

In each training menu, we counted the number of training sets performed by each participant up to the day before the second MRI day. To evaluate between-group differences in the number of training sets performed during the training period, we performed a two-sample *t*-test for each menu. We found no significant difference in any of the menus (Table S3). In addition, we confirmed the absence of significant between-group differences in the mean number of sets of each training menu performed on the day before the second MRI day. Finally, as for the subgroups of the BM group, we also confirmed the absence of significant differences between the two subgroups ($t[21] = 1.54$, $P = 0.14$).

**Kinesthetic illusion task**

Before the fMRI experiment, each participant experienced this task outside the MRI room, to familiarize them with the task. Participants then entered the room and were placed in the MRI scanner. Their heads were immobilized using sponge cushions and adhesive tape, and their ears were plugged. Both arms were naturally semi-pronated, extended along their body, and were supported by cushions, thus allowing the participants to relax their upper arms during the task. Each participant completed two experimental runs.

After each experimental run, we asked the participants whether they felt the illusion during tendon vibration and bone vibration. To verify that the participants experienced a substantial kinesthetic illusion of right-hand flexion during the tendon-vibration task, we asked them to remember the maximum illusory flexion angle experienced during a run and to indicate the maximum angle after the run. In the scanner (immediately after each run), we showed the participants a protractor on which a hand-shape indicator was mounted (they could see the protractor through a mirror). Before the participants entered the scanner room, we measured the flexion angle of the relaxed right wrist of each participant using a protractor. The hand-shape indicator was first set at this angle, which corresponded to the original position of the relaxed hand of each participant. From this position, we began flexing the indicator. The participants were instructed to say “stop” when they believed that the indicator had reached the maximum illusory angle that they had experienced. We measured this angle as the change from the original position. We defined this as the illusory angle, then calculated the mean illusory angle between two runs for each participant. We first evaluated the possible between-age-group (younger, older) difference using a two-sample *t*-test. To evaluate the training effect on the illusory angle, we conducted a two-way ANOVA that included one between-subject factor (group [2]: BM or RH) and one within-subject factor (order [2]: before or after).

**MRI data acquisition**

Functional images were acquired using T2*-weighted gradient echo-planar imaging (EPI) sequences on a 3.0-Tesla MRI scanner (Trio Tim; Siemens, Germany) equipped with a 32-

channel array coil. Each volume consisted of 44 slices (slice thickness, 3.0 mm; inter-slice thickness, 0.5 mm) acquired in ascending order, covering the entire brain. The time interval between successive acquisitions from the same slice was 2,500 ms. An echo time of 30 ms and a flip angle of 80° were used. The field of view was 192 × 192 mm, and the matrix size was 64 × 64 pixels. The voxel dimensions were 3 × 3 × 3.5 mm in the x-, y-, and z-axes, respectively. We collected 65 volumes for each experimental run. As an anatomical reference, a T1-weighted magnetization-prepared rapid gradient echo (MP-RAGE) image was acquired using the same scanner. The imaging parameters were as follows: TR = 1900 ms, TE = 2.48 ms, FA = 9°, field-of-view (FOV) = 256 × 256 $mm^2$, matrix size = 256 × 256 pixels, slice thickness = 1.0 mm, voxel size = 1 × 1 × 1 $mm^3$, and 208 contiguous transverse slices. The same MRI data acquisition was performed before and after training.

**fMRI data preprocessing**

To eliminate the effects of unsteady magnetization during the tasks, we discarded the first four EPI images in each fMRI run before the start of the first epoch. Imaging data were analyzed using SPM 12 (default setting: Wellcome Trust Centre for Neuroimaging, London, UK) implemented in MATLAB (MathWorks, Sherborn, MA, USA). To compare EPI images before and after training, we used the T1-weighted image acquired before the training as the anatomical target image. EPI images were aligned to the first image. Through this realignment procedure, we obtained head-position data, which changed over time from the first frame through six parameters. All participants had a maximum displacement of < 1.5 mm in the x, y, or z-plane and an angular rotation about each axis < 0.1° during an fMRI run. Thus, no data were excluded from the analysis. The realigned EPI images were co-registered to the T1-weighted structural image of each participant that was acquired before the training and was spatially normalized to the standard stereotactic Montreal Neurological Institute (MNI) space [3] with 2-mm isotropic voxel size using the SPM12 normalization algorithm. Finally, the normalized images were filtered using a Gaussian kernel with a full-width at half-maximum of 4 mm along the x-, y-, and z-axes.

### Supplementary Results

**Temporal profile of task-related brain activity during the illusion in the contralateral (left) and ipsilateral (right) SM1-PMD regions before training in the younger and older groups**

To assess the temporal profile of task-related brain activity during the illusion in the contralateral (left) and ipsilateral (right) hand/arm section of the SM1 and PMD regions before training in the younger and older groups carefully, we performed an additional analysis. For this analysis, we prepared ipsilateral (right) SM1-PMD and contralateral (left) ROIs, which were also defined based on the active cluster identified during the left-hand and right-hand illusion in our previous study

[4].

We extracted the time-course data of brain activity from 18 volumes immediately before, during, and immediately after each tendon-vibration epoch (six volumes during the pre-tendon-vibration resting period, six volumes during the tendon-vibration epoch, and six volumes during the post-tendon-vibration resting period). This was carried out for each of the first, second, third, and fourth tendon-vibration epochs in each run for each participant (the fifth tendon-vibration epoch in each run was excluded because of the lack of a post-tendon-vibration resting period). The time-course data were extracted from each of the contralateral (left) and ipsilateral (right) SM1-PMD ROIs defined in our previous study (Figure S1a). We normalized the time-course data of each voxel by subtracting its temporal mean and dividing it by its standard deviation in each run. We then averaged the data obtained from all voxels in each of the contralateral and ipsilateral SM1-PMD ROIs for each participant. Finally, we averaged the normalized time-course data related to the eight tendon-vibration epochs for each participant and calculated the grand average across all participants in each age group (n = 31 for the younger participants, n = 48 for the older participants). When analyzing the grand average data obtained from the bilateral SM1-PMD ROIs in each age group, we found that the ipsilateral SM1-PMD activity was suppressed during tendon vibration in the younger group, but not in the older group, although the contralateral SM1-PMD activity was increased in both age groups (Figure S1b, c).

**Additional analysis of the relationship between deteriorated dexterity and ipsilateral M1 activity in older adults before training**

Even though we formulated a strong anatomical hypothesis regarding the correlation between deteriorated dexterity and ipsilateral M1 activity in older adults, one might argue that the cluster size (6 voxels) was too small. To examine the fortification of the ipsilateral M1 cluster with the increase in the number of participants, we recruited 24 additional healthy, right-handed, older adults (age, 71.7 ± 3.1 years; males, 15) from the same local human resources center for older adults. We assessed their cognitive status using the Mini-Mental State Examination (MMSE). Their average MMSE score was 28.5 ± 1.8, which was not significantly different from that obtained for the BM and RH groups (see main text). We also confirmed the handedness of participants using the Edinburgh Handedness Inventory [5]. No participants had a history of neurological, psychiatric, or movement disorders, as assessed based on self-reports. We explained the details of the MRI experiment before the experiment, after which participants provided written informed consent. The participants completed the same peg task, and we measured their brain activity during the right-hand illusion using the same scanner and scanning parameters, and analyzed their data in the same way as in the present study.

The average peg time across the 24 older adults was 21.66 ± 3.40 s (mean ± standard deviation), which was highly similar to that obtained from the RH (22.04 ± 3.78 s) and BM (22.23

± 3.58 s) groups. We examined brain regions in which the activity was positively correlated with the peg time (height-threshold $P < 0.005$, uncorrected) in the 72 older adults, as we did for the 48 older participants in the present study. We found the largest cluster showing the highest peak value in the hand/arm section of the ipsilateral M1 in the whole brain (67 voxels, peak voxel = 38, −28, 64; T = 4.91, cytoarchitectonic area 4a) (Figure S2a). This cluster overlapped with the M1 ROI; thus, we found a significant cluster of voxels when we evaluated its spatial extent within the M1 ROI (extent-threshold $P < 0.05$, FWE-corrected). The peak (38, −28, 64) was highly similar to that observed in the analysis of the 48 participants (36, −28, 66). Thus, the ipsilateral M1 cluster observed in the 48 participants became robust when we increased the number of participants. We visualized the relationship between the activity obtained from the M1 ROI and the peg time across participants and confirmed a higher positive correlation ($r = 0.49$, n = 72; Figure S2b). This series of results strengthened the present findings of the correlation between the reduction/loss of ipsilateral M1 deactivation and deteriorated dexterity in older adults.

**Analysis of training effect on functional connectivity**

Because the BM group underwent bimanual training, we expected that the training would change the functional connectivity between the activities in the hand/arm sections of the bilateral sensorimotor regions (SM1-PMD) through interhemispheric interaction. Therefore, we examined possible between-training-group differences (RH vs. BM and BM vs. RH) in the change in functional connectivity with the hand/arm section of the contralateral (left) sensorimotor regions after the training. We performed a PPI analysis [6,7], as implemented in the Generalized PPI toolbox (https://www.nitrc.org/projects/gppi). We used the contralateral (left) SM1-PMD ROI (see above) as a seed region in the analysis. First, for each participant, the time course of the average fMRI signal across the voxels in the contralateral (left) SM1-PMD ROI was deconvolved using the canonical hemodynamic response function (physiological variable). Next, we performed a general linear model analysis using the design matrix and including the following regressors: the physiological variable, boxcar function for the task epoch (psychological variable), and multiplication of the physiological variable and the psychological variable (PPI). These variables were convolved with a canonical hemodynamic response function. Six realignment parameters were also included in the design matrix as regressors of no interest.

We generated images of voxels in which activity changed with the PPI regressor in each participant, to depict voxels in which activity changed in association with the fluctuation of the left SM1-PMD activity during the task epoch. We generated contrast images (after–before) for each participant, which were used in the second-level group analysis, in which we examined possible between-training-group differences in the change in functional connectivity with the hand/arm section of the contralateral (left) SM1-PMD after the training. We examined significant clusters throughout the brain. We also carefully checked for possible between-training-group differences in the ipsilateral SM1-PMD ROI. In this analysis, we extracted the effect size of the connectivity change from the ROI in each participant and calculated the average effect size across participants in each training group. We used a two-sample *t*-test to evaluate the between-training-group differences.

First, we examined brain regions in which the connectivity decreased during the illusion after the training in the BM group compared with the RH group using the contrast of RH vs. BM. We found a significant cluster in the ipsilateral SM1-PMD region (pink and white areas in Figure S3a), in addition to the contralateral parietal and visual cortices, in the entire brain (see Table S8). Most of the voxels (103 out of 113 voxels) in the ipsilateral cluster were located within the ipsilateral SM1-PMD ROI (white area in Figure S3a). In contrast, we found no clusters that showed significant differences in the opposite contrast of BM vs. RH in the entire brain.

Regarding the ipsilateral SM1-PMD ROI, we analyzed the individual effect size of connectivity in this ROI in each group. We found a significant between-group difference ($t$[46] = 2.96, $P$ = 0.005; Figure S3b), indicating that the interhemispheric connectivity between two SM1-PMD ROIs was decreased in the BM group compared with the RH group. A different training effect on functional connectivity was observed with no significant interaction (group [BM and RH] × order [before and after]) in the illusory angle (see Table S6).

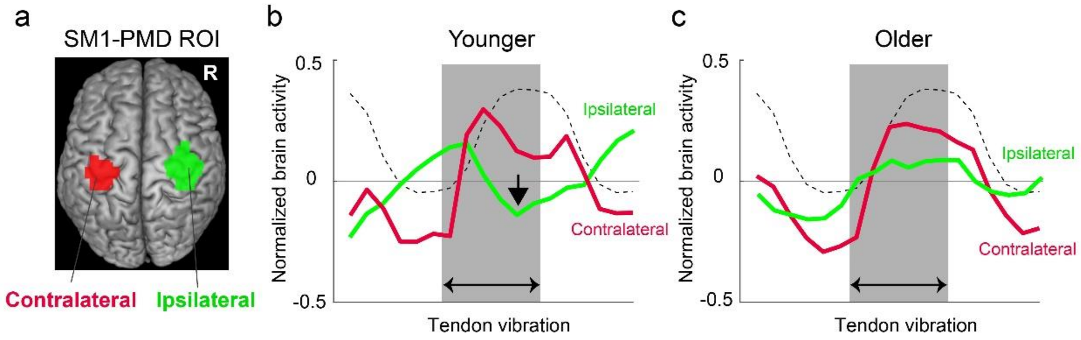


**Fig. S1.** Temporal profile of brain activity (hemodynamic response) related to the tendon-vibration (illusion) before training in the contralateral (left) and ipsilateral (right) SM1-PMD ROIs for the younger and older groups. a: Contralateral (red) and ipsilateral (green) SM1-PMD ROIs. b: Grand average data obtained from the younger group. c: Grand average data obtained from the older group. b, c: In each panel, the red line indicates the grand average of brain activity across participants obtained from the contralateral ROI. The green line indicates the grand activity obtained from the ipsilateral ROI, and the dashed curved line indicates a boxcar function convolved with a canonical hemodynamic response function. The gray period represents the tendon-vibration epoch. The vertical axis indicates the level of normalized brain activity. The activity in the contralateral SM1-PMD ROI increased during the tendon-vibration epoch in both the younger and older groups. However, the activity in the ipsilateral SM1-PMD ROI was suppressed in the younger adults (downward arrow), but not in the older adults (C). Abbreviations: R, right hemisphere

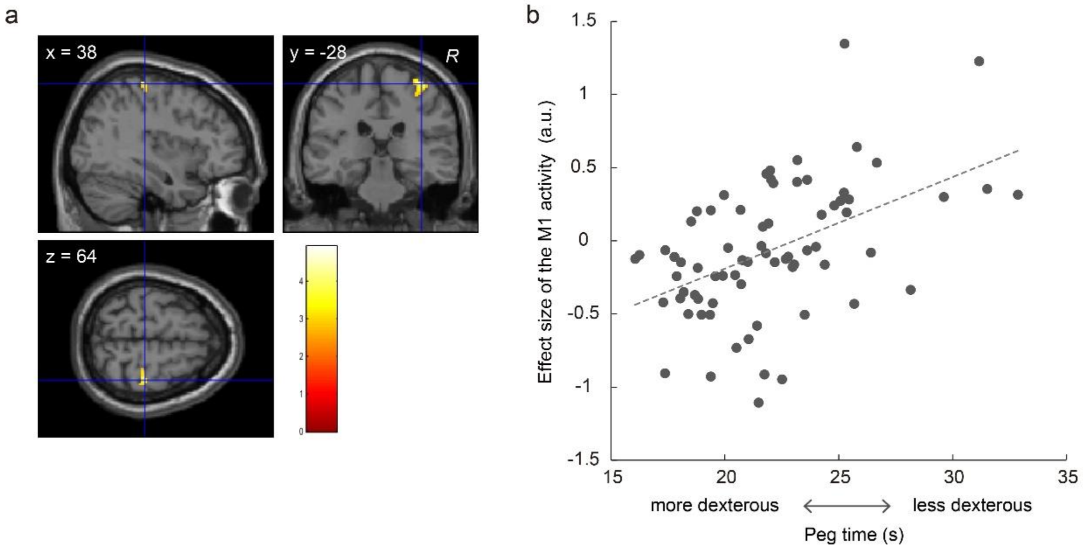


**Fig. S2.** a: The ipsilateral M1 region, in which the activity was positively correlated with the peg time in 72 older adults. This cluster (67 voxels) was observed in the highly similar ipsilateral M1 region observed in the analysis of the 48 participants. The crossing point between the two blue lines indicates a peak voxel (38, −28, 64) in the ipsilateral M1 (cytoarchitectonic area 4a). b: Relationship between the effect size of activity in the M1 ROI (a.u.; vertical axis) and the peg time (horizontal axis) across 72 older adults. Each dot represents an individual datum, and the dashed line indicates a regression line fitted to the data ($r$ = 0.49, n = 72). Abbreviations: a.u., arbitrary unit; R, right hemisphere

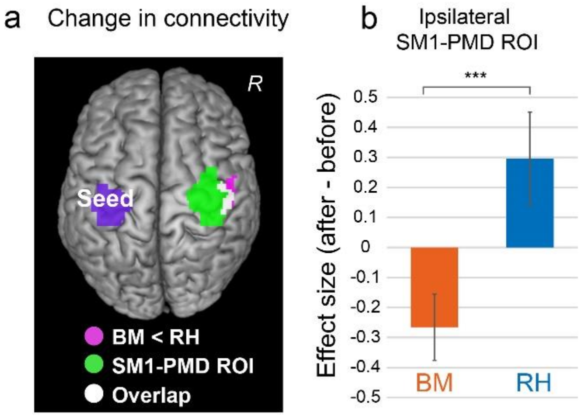


**Figure S3.** Training effects on functional connectivity during the illusion. a: Brain regions (pink and white areas) that showed significant between-training-group differences (RH > BM) in changes in functional connectivity during the illusion after the training. The purple area indicates the seed region (i.e., contralateral SM1-PMD ROI), and the green and white areas indicate ipsilateral SM1-PMD ROIs. The white area indicates an overlapping between the region that showed the group difference and the ipsilateral SM1-PMD ROI. b: Average effect size of functional connectivity changes (a.u.; vertical axis) in the ipsilateral SM1-PMD ROI (after–before) across participants in each (BM, RH) group. The error bars indicate the SEM. *** $P \leq 0.005$. Abbreviations: BM, bimanual; RH, right hand. Abbreviations: BM, bimanual; R, right hemisphere; RH, right hand; SEM, standard error of the mean

**Table S1**. Training menus

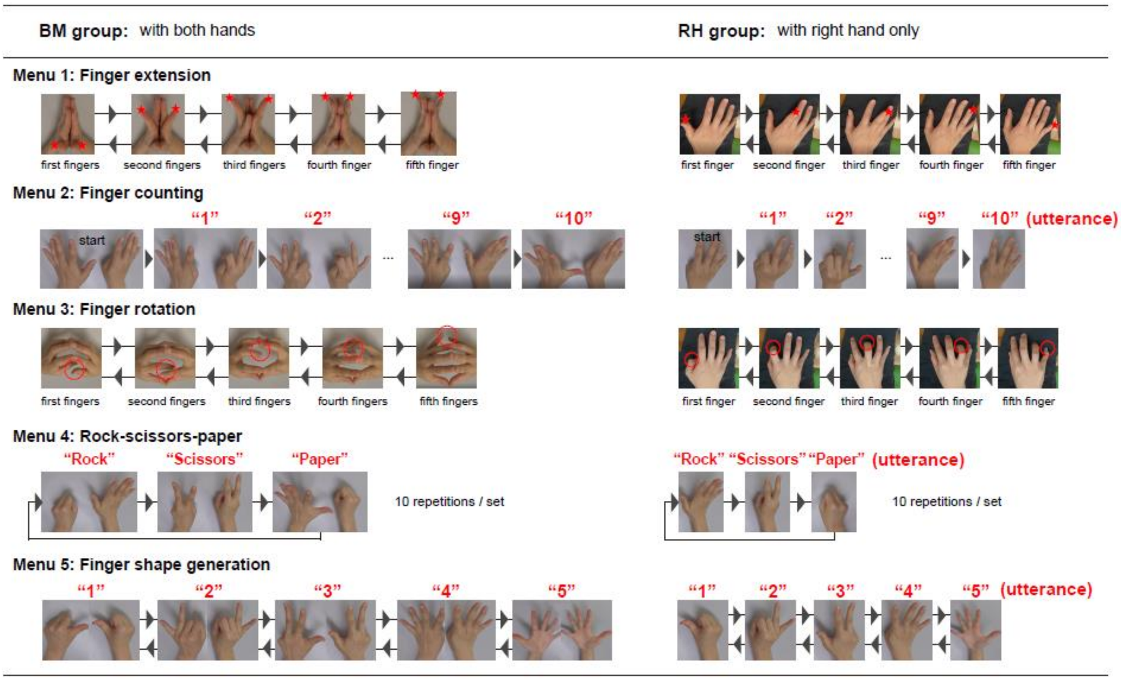

**Table S2.** Homework menus (sets that needed to be performed each week)

| | Homework menus |
|---|---|
| Week 1 | Menus 1 (3 sets) and 2 (3 sets) |
| Week 2 | Menus 1 (3 sets), 2 (3 sets), and 3 (3 sets) |
| Week 3 | Menus 1 (3 sets), 2 (3 sets), and 3 (3 sets) |
| Week 4 | Menus 1 (5 sets), 2 (5 sets), and 3 (5 sets) |
| Week 5 | Menus 1 (5 sets), 3 (5 sets), and 4 (3 sets) |
| Week 6 | Menus 1 (5 sets), 3 (5 sets), and 4 (3 sets) |
| Week 7 | Menus 3 (5 sets), 4 (3 sets), and 5 (3 sets) |
| Week 8 | Menus 3 (5 sets), 4 (5 sets), and 5 (5 sets) |

1: Finger extension; 2: Finger counting; 3: Finger rotation; 4: Rock–paper–scissors; 5: Finger shape generation

**Table S3.** Mean number of sets of each training menu performed by each group

| | **BM group** (n = 23) | | **RH group** (n = 25) | | |
|---|---|---|---|---|---|
| Menu | Mean | SD | Mean | SD | *P*-value |
| 1. Finger extension | 230.3 | 149.7 | 240.4 | 135.2 | 0.81 |
| 2. Finger counting | 149.8 | 140.0 | 131.3 | 96.7 | 0.60 |
| 3. Finger rotation | 208.6 | 135.0 | 211.1 | 71.5 | 0.94 |
| 4. Rock–paper–scissors | 101.0 | 31.1 | 100.2 | 47.6 | 0.95 |
| 5. Finger shape generation | 44.2 | 18.8 | 52.4 | 22.9 | 0.20 |
| Total | 733.9 | 427.6 | 735.5 | 305.1 | 0.99 |

**Table S4.** Brain regions showing task-related deactivations and activations in the younger and older groups

| | Size | Cluster-level Corrected *P* | x | y | z | T-value | Area |
|---|---|---|---|---|---|---|---|
| **Older group** | | | | | | | |
| (a) Brain deactivations before training | | | | | | | |
| | 1076 | < 0.001 | 48 | −72 | 30 | 6.47 | Angular gyrus (Area PGp) |
| | | | −56 | −64 | 10 | 6.17 | Middle temporal gyrus (Area PGp) |
| | | | −64 | −42 | 10 | 6.06 | Middle temporal gyrus (Area TE3) |
| | 981 | < 0.001 | 62 | −44 | 8 | 6.10 | Middle temporal gyrus (Area PGa) |
| | | | 56 | −42 | 14 | 5.46 | Superior temporal gyrus (Area PFm) |
| | | | 50 | −52 | 20 | 5.37 | Middle temporal gyrus (Area PGa) |
| | 301 | < 0.001 | −62 | −14 | 18 | 6.08 | Postcentral gyrus (Area OP4) |
| | | | −50 | −18 | 20 | 5.03 | Postcentral gyrus (Area OP1) |
| | | | −58 | −16 | 30 | 3.97 | Postcentral gyrus (Area 3b) |
| | 1685 | < 0.001 | 4 | −62 | 34 | 5.62 | Precuneus |
| | | | 4 | −58 | 46 | 5.17 | Precuneus |
| | | | 10 | −48 | 38 | 5.10 | Precuneus |
| | 261 | < 0.001 | 4 | 44 | −6 | 4.95 | Middle Orbital gyrus (Area s32) |
| | | | 8 | 58 | −2 | 4.08 | Middle Orbital gyrus (Area Fp2) |
| | | | 4 | 54 | −8 | 3.79 | Middle Orbital gyrus (Area Fp2) |
| | 161 | < 0.05 | −54 | −8 | 44 | 4.32 | Postcentral gyrus |
| | | | −50 | −10 | 30 | 4.05 | Postcentral gyrus (Area 4p) |
| | | | −50 | −18 | 38 | 3.36 | Postcentral gyrus (Area 3b) |
| (b) Brain activations before training | | | | | | | |
| | 17106 | < 0.001 | −32 | −26 | 52 | 11.99 | Postcentral gyrus (Area 4p) |
| | | | 56 | 10 | 14 | 8.60 | Inferior frontal gyrus (Area 44) |
| | | | 50 | 12 | 2 | 8.28 | Inferior frontal gyrus (Area 44) |
| | 4303 | < 0.001 | 18 | −46 | −20 | 9.76 | Lobule V (Hem) |
| | | | 26 | −56 | −52 | 8.32 | Lobule VIIIa (Hem) |
| | | | 14 | −54 | −14 | 7.55 | Lobule VI (Hem) |

| | | | | | | |
|---|---|---|---|---|---|---|
| 2228 | < 0.001 | −34 | 18 | 2 | 7.23 | Insula lobe |
| 544 | < 0.001 | 62 | −30 | 40 | 5.83 | Supramarginal gyrus (Area PF) |
| | | 60 | −22 | 40 | 4.88 | Supramarginal gyrus (Area PFt) |
| | | 62 | −28 | 30 | 4.47 | Supramarginal gyrus (Area PF) |
| 313 | < 0.001 | −32 | −60 | −54 | 4.92 | Lobule VIIIa (Hem) |
| | | −26 | −68 | −54 | 4.82 | Lobule VIIb (Hem) |
| | | −26 | −44 | −48 | 4.64 | Lobule VIIIb (Hem) |
| **Younger group** | | | | | | |
| (c) Brain deactivations | | | | | | |
| 10140 | < 0.001 | 40 | −20 | 66 | 8.82 | Precentral gyrus |
| | | 0 | −52 | 38 | 8.56 | Precuneus |
| | | 48 | −18 | 58 | 8.38 | Postcentral gyrus |
| 2869 | < 0.001 | 54 | −44 | 14 | 8.72 | Superior temporal gyrus (Area PFm) |
| | | 64 | −32 | 10 | 7.15 | Superior temporal gyrus (Area TE3) |
| | | 38 | −14 | 20 | 6.76 | Insula lobe (Area OP3) |
| 4690 | < 0.001 | −54 | −12 | 12 | 8.08 | Superior temporal gyrus (Area OP4) |
| | | −60 | −2 | 18 | 7.30 | Postcentral gyrus (Area 3b) |
| | | −50 | −14 | 44 | 7.14 | Postcentral gyrus (Area 4a) |
| 2819 | < 0.001 | 2 | 38 | −4 | 6.27 | ACC (Area s24) |
| | | 2 | 50 | −10 | 6.22 | Middle orbital gyrus (Area Fp2) |
| | | 10 | 42 | −12 | 5.87 | Middle orbital gyrus (Area s32) |
| 734 | < 0.001 | −22 | 30 | 52 | 5.69 | Middle frontal gyrus |
| | | −24 | 22 | 40 | 5.21 | Middle frontal gyrus |
| | | −14 | 42 | 36 | 4.93 | Superior frontal gyrus |
| (d) Brain activations | | | | | | |
| 4835 | < 0.001 | −30 | −26 | 54 | 10.33 | Precentral gyrus (Area 4p) |
| | | −2 | −10 | 58 | 8.19 | Posterior-medial frontal |
| | | 2 | 10 | 52 | 7.35 | Posterior-medial frontal |
| 1777 | < 0.001 | 18 | −48 | −18 | 8.40 | Lobule V (Hem) |
| | | 36 | −54 | −26 | 7.08 | Lobule VI (Hem) |

| | | | | | | |
|---|---|---|---|---|---|---|
| | | 26 | −44 | −24 | 6.50 | Lobule VI (Hem) |
| 10422 | < 0.001 | 32 | 18 | 10 | 8.08 | Insula lobe |
| | | −30 | 22 | 8 | 8.04 | Insula lobe |
| | | 54 | 8 | 24 | 7.77 | Inferior frontal gyrus (Area 44) |
| 1467 | < 0.001 | 60 | −22 | 36 | 6.11 | Supramarginal gyrus (Area PFt) |
| | | 36 | −32 | 44 | 6.09 | Postcentral gyrus (Area 3a) |
| | | 46 | −38 | 48 | 5.89 | Inferior parietal lobule (Area hIP2) |
| 425 | < 0.001 | −32 | −52 | −26 | 6.07 | Lobule VI (Hem) |
| | | −24 | −68 | −20 | 4.93 | Lobule VI (Hem) |
| | | −40 | −66 | −26 | 4.54 | Lobule VIIa crusI |
| 249 | < 0.001 | −28 | −58 | −54 | 4.57 | Lobule VIIIa (Hem) |
| | | −20 | −70 | −46 | 4.54 | Lobule VIIb (Hem) |
| | | −20 | −64 | −52 | 4.09 | Lobule VIIIa (Hem) |

ACC: anterior cingulate cortex

**Table S5.** Brain regions showing between-age-group differences

| Size | Cluster-level Corrected *P* | X | Y | z | T-value | Area |
|---|---|---|---|---|---|---|
| **(a) Older vs. younger** | | | | | | |
| 3521 | < 0.001 | 28 | −24 | 54 | 7.10 | Precentral gyrus (Area 4p) |
| | | 36 | −22 | 48 | 6.70 | Postcentral gyrus (Area 4p) |
| | | 40 | −20 | 66 | 6.50 | Precentral gyrus |
| 1849 | < 0.001 | −2 | −50 | 38 | 5.89 | Precuneus |
| | | 14 | −68 | 24 | 4.84 | Cuneus |
| | | 0 | −44 | 46 | 4.72 | MCC |
| 407 | < 0.001 | −44 | −30 | 10 | 4.93 | Superior temporal gyrus (Area TE 1.1) |
| | | −50 | −54 | 18 | 4.46 | Middle temporal gyrus (Area PGa) |
| | | −34 | −32 | 20 | 4.23 | Insula lobe (Area OP2) |
| 433 | < 0.001 | 64 | −30 | 10 | 4.88 | Superior temporal gyrus (Area TE 3) |
| | | 54 | −44 | 14 | 4.34 | Superior temporal gyrus (Area PFm) |
| | | 44 | −54 | 10 | 3.93 | Middle temporal gyrus |
| 384 | < 0.001 | −50 | −14 | 44 | 4.86 | Postcentral gyrus (Area 4a) |
| | | −60 | −2 | 18 | 4.79 | Postcentral gyrus (Area 3b) |
| | | −58 | −6 | 26 | 4.16 | Postcentral gyrus (Area 3b) |
| 197 | < 0.005 | 0 | 34 | 6 | 4.52 | ACC (Area 33) |
| | | 4 | 40 | 0 | 3.86 | ACC (Area s24) |
| | | 0 | 56 | 8 | 3.52 | Superior medial gyrus (Area Fp2) |
| 204 | < 0.005 | −12 | −84 | 28 | 4.05 | Cuneus (Area hOc3d) |
| | | −20 | −82 | 26 | 3.98 | Superior occipital gyrus |
| | | −40 | −78 | 18 | 3.20 | Middle occipital gyrus (Area PGp) |
| **(b) younger vs. older** | | | | | | |
| 188 | < 0.01 | −16 | −22 | 18 | 4.76 | Thalamus (prefrontal) |
| | | −24 | −16 | 14 | 4.25 | Thalamus (motor) |
| | | −20 | −12 | 22 | 3.91 | Caudate nucleus |

ACC: anterior cingulate cortex; MCC: middle cingulate cortex

**Table S6.** Illusory angles in younger and older (BM and RH) groups (unit: degree)

| | **Younger group** | **Older group** | | | |
|---|---|---|---|---|---|
| | (n = 31) | BM group (n = 23) | | RH group (n = 25) | |
| | | Before | After | Before | After |
| Mean | 32.82 | 35.38 | 37.83 | 29.85 | 31.70 |
| SD | 13.37 | 17.16 | 16.08 | 14.66 | 11.00 |

Note: There was no significant interaction ([BM and RH] × [before and after]) in the illusory angle of the older group ($F[1, 46] = 2.21$, n.s.).

**Table S7.** Brain regions in which the activity reduction was correlated with the dexterity improvement in the BM group

| Cluster-level | | | | | | |
|---|---|---|---|---|---|---|
| Size | Corrected *P* | X | y | Z | T-value | Area |
| 114 | < 0.05 | −2 | −12 | 42 | 7.53 | MCC |
| | | 2 | −24 | 44 | 5.65 | MCC |
| | | −4 | −20 | 36 | 4.01 | MCC |
| 109 | < 0.05 | 22 | −28 | 74 | 5.67 | Precentral gyrus (Area 4a) |
| | | 20 | −30 | 64 | 4.51 | Precentral gyrus (Area 4p) |
| | | 32 | −28 | 70 | 4.20 | Precentral gyrus (Area 4a) |

MCC: middle cingulate cortex

**Table S8.** Training effect on functional connectivity: $(\text{after–before})_{RH}$ – $(\text{after–before})_{BM}$

| Cluster-level | | | | | | |
|---|---|---|---|---|---|---|
| Size | Corrected *P* | X | Y | z | T-value | Area |
| 113 | < 0.05 | 48 | −22 | 48 | 5.35 | Postcentral gyrus (Area 2) |
| | | 48 | −10 | 56 | 4.02 | Precentral gyrus (Area 4a) |
| | | 54 | −14 | 48 | 3.45 | Postcentral gyrus (Area 1) |
| 135 | < 0.01 | −34 | −46 | 54 | 5.07 | Inferior parietal lobule (Area 7PC) |
| | | −48 | −44 | 48 | 3.96 | Inferior parietal lobule (Area hIP2) |

| | | | | | | |
|---|---|---|---|---|---|---|
| | | −40 | −48 | 62 | 3.92 | Superior parietal lobule (Area 7PC) |
| 138 | 0.005 | −6 | −82 | 8 | 4.31 | Calcarine gyrus (Area V1) |
| | | −16 | −68 | 4 | 4.27 | Calcarine gyrus (Area V1) |
| | | −10 | −56 | 2 | 4.11 | Lingual gyrus (Area V1) |
